# Gelsolin regulates proliferation, apoptosis and invasion in NK/T-cell lymphoma cells through the PI3K/Akt signaling pathway

**DOI:** 10.1101/152603

**Authors:** Yanwei Guo, Hongqiao Zhang, Xin Xing, Lijuan Wang, Jian Zhang, Lin Yan, Xiaoke Zheng, Mingzhi Zhang

## Abstract

The expression of gelsolin (GSN) is abnormal in many cancers, including extranodal nasal-type natural killer/T-cell lymphoma (NKTCL). However, the biological function of GSN and its mechanism in NKTCL remain unclear. We found GSN overexpression significantly suppressed cell proliferation, colony formationand invasion and promoted apoptosis of YTS cells. Moreover, the upregulation of GSN significantly decreased the protein levels of PI3K and p-AKT. Interestingly, blocking the PI3K/AKT signaling pathway significantly inhibited cell proliferation and invasion and promoted apoptosis of YTS cells. In conclusion, our findings indicate that GSN can suppress cell proliferation and invasion and promote apoptosis of YTS cells, which is likely to be mediated at least partially through inhibition of the PI3K/AKT signaling pathway.

## Introduction

Extranodal nasal-type natural killer/T-cell lymphoma (NKTCL) is one of the EBV related hematological malignancies, which mainly develops in the nasal cavity but can also occur in extranasal sites, either as a primary extranasal or disseminated disease (Harabuchi et al., 1996, Chen et al., 2015). NKTCL is more common in Asia than in Western countries (Au et al., 2009). Although most of the cases of NKTCL are diagnosed in the early stage of the disease, the long term survival rate of patients is about 46%-60% (Suzuki et al., 2010). The one year survival rate of the patients with advanced-stage disease is only 50%, despite improvements in treatment (Jaccard et al., 2011, Yamaguchi et al., 2011). The tumor cells of NKTCL derived from NK cells and rarely T cell is linked to Epstein-Barr virus (EBV) infection (Huang et al., 2013). However, the biological characteristics of NKTCL are not yet to be completely clear.

Gelsolin (GSN), as a Ca^2+^-regulated actin filament severing and capping, is a widespread, polyfunctional regulator of cell structure and metabolism (Li et al., 2012). GSN is a widely expressed actin regulator, and has been reported to be a multifunctional regulator of physiological and pathological cellular process and regulate cell migration, cell morphology, proliferation and apoptosis (Sun et al., 1999, Li *et al*. 2012). Previous research demonstrated that GSN was prevalently expressed in a variety of cells (Tanaka et al., 2006). A previous study revealed that the levels of GSN are decreased in various cancer, including breast, urinary bladder, colon, kidney, ovary, prostate, gastric and urinary system cancer (Tanaka *et al*. 2006). A study presented by Zhou *et al*. showed that up-regulated GSN inhibits apoptosis while down-regulated GSN promotes apoptosis, which could be associated with the regulation of GSN on the apoptosis associated pathways and the apoptosis factors caspase 3 and bcl-2 (Zhou et al., 2015). In addition, a study showed that GSN was observed *in vitro* to suppress the proliferation and invasion of 786-0 renal cell carcinoma cells (Zhu et al., 2015). A previous study found that GSN in colorectal tumor cell regulates cell invasion through its modulation of the uPA/uPAR cascade, with possible vital roles in colorectal tumor dissemination to metastatic sites (Zhuo et al., 2012).

GSN displayed high expression in the secondary diffuse large B-cell lymphoma (DLBCL) compared with de novo DLBCL (Ludvigsen et al., 2015). However, a recent study revealed that the level of GSN is downregulated in serums of the advanced NKTCL patients (Zhou et al., 2016). Despite the roles of GSN has been explored, whether the GSN could modulate cell proliferation, apoptosis and invasion in NK/T-cell lymphoma cells is currently unknown. Further investigations are required concerning the role of GSN in NK/T-cell lymphoma progression to explore whether decreased or increased GSN level in NK/T-cell lymphoma exists a direct relationship with tumorigenesis.

Here, to investigate the effects of GSN on the proliferation, apoptosis, and invasion of NK/T-cell lymphoma cells and the potential molecular mechanism, GSN was overexpressed in YTS cell line *in vitro*, which may contribute to the current understanding of the biological functions of GSN in NK/T-cell lymphoma.

## Results

### Overexpression of GSN in transfected YTS cells

After the Lenti-virus containing Lenti-GSN vector was transfected into YTS cells, green fluorescence was obvious in the infected YTS cells, as observed under a fluorescence microscope, and the result indicated a successful transfection (Fig. 1 A). Flow cytometry analysis showed that the transfection ratio in cells was 70-80% (Fig. 1 B). qRT-PCR analysis and western blot analysis demonstrated that the mRNA and protein levels of GSN were both significantly increased in the YTS-GSN cells, when compared with the YTS-Con cells (Fig. 1C and D).

**Figure 1.**
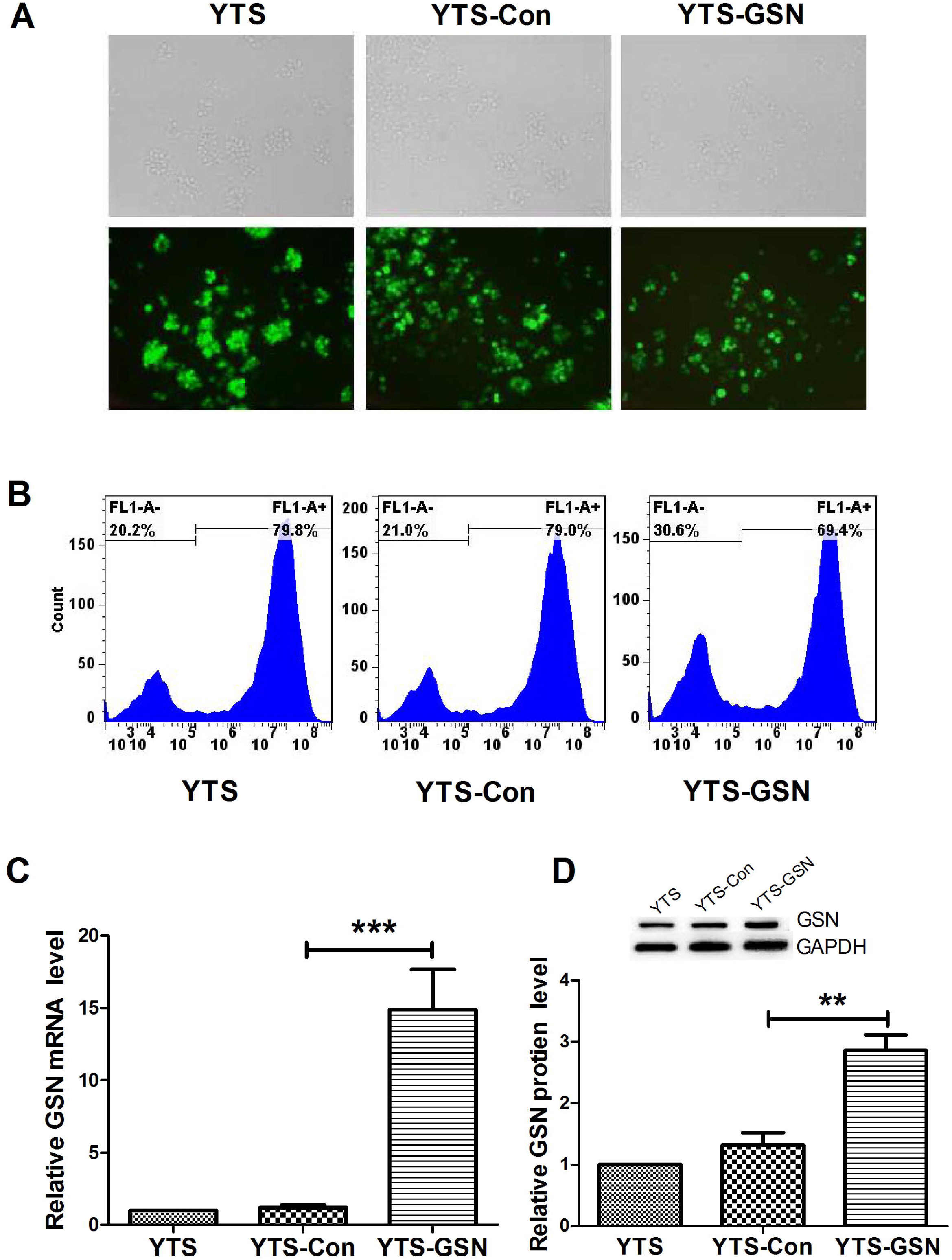
Transfection ratio of the Lenti-virus containing Lenti-GSN vector and GSN overexpression in YTS cells. (A) Green fluorescence was observed in the transfected YTS cells under a fluorescence microscope (magnification, ×200) at 48 h, which indicated a successful transfection. (B) Flow cytometric analysis demonstrated that the transfection ratio in cells was 70-80% at 48 h. (C) qRT-PCR analysis exhibited that GSN mRNA expression was higher in YTS-GSN cells than that in YTS- Con cells at 48 h. (D) Western blot analysis showed that the protein level of GSN was higher in YTS-GSN cells than that in YTS- Con cells at 48 h. GSN: gelsolin; YTS cells: no transfected cells; YTS-Con cells: YTS cells transfected with the pCDH-CMV-MCS-EF1-copGFP vector; YTS-GSN cells: YTS cells transfected with the pCDH-CMV-MCS-EF1-copGFP-GSN vector. \*\**P* < 0.01, \*\*\**P* < 0.001.

### GSN overexpression inhibits YTS cell proliferation and colony formation

To explore the effects of GSN on YTS cell proliferation and colony formation, CCK-8 assay and colony formation assay were performed. Our results of CCK-8 assay revealed that cell proliferation of YTS-GSN cells was significantly suppressed, compared with YTS-Con cells, respectively (Fig. 2A). In addition, the results of colony formation assay demonstrated that GSN resulted in a decrease in the clonogenic survival of YTS-GSN cells, compared with YTS-Con cells (Fig. 2B). These results suggested GSN had inhibitory effect on YTS cell proliferation.

**Figure 2.**
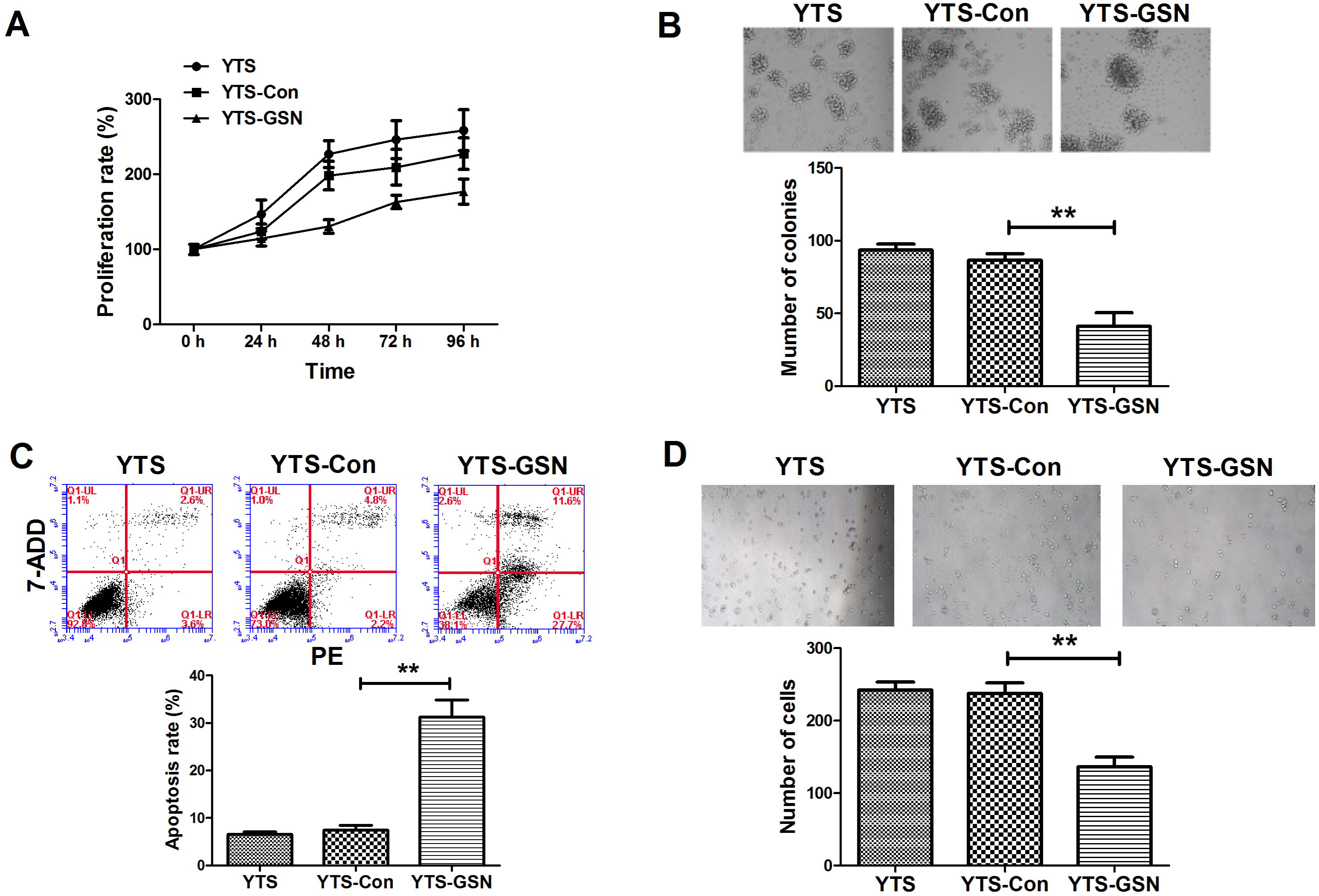
GSN overexpression inhibits proliferation, colony formation and invasion of YTS cells and promotes apoptosis. (A) CCK-8 assays revealed that GSN overexpression inhibited YTS cell proliferation. (B) Shown were representative photomicrographs of colony formation assay of YTS cells transfected with lenti-Con and lenti-GSN plasmids for 12 days. Statistical analysis of colony formation assay showed that GSN overexpression caused a decrease in the clonogenic survival of YTS cells compared with YTS- Con cells. (C) The representative photomicrographs of flow cytometric analysis were shown. Statistical analysis of flow cytometric analysis showed that GSN overexpression significantly increased apoptosis rate of YTS cells at 48 h. (D) Representative photomicrographs of transwell invasion assay in cells at 48 h and statistical analysis showed that GSN obviously suppressed YTS cell invasion. \*\**P* < 0.01.

### GSN overexpression promotes apoptosis and inhibits YTS cell invasion

Next, we detected the roles of GSN on apoptosis and YTS cell invasion. Flow cytometry analysis revealed that GSN overexpression caused a significant increase in apoptotic cell of YTS cells (Fig. 2C). Transwell assay showed that invasion of YTS-GSN cells was significantly inhibited, compared with YTS-Con cells (Fig. 2D).

### GSN overexpression suppresses the PI3K/Akt pathway in YTS cells

To further confirm the potential mechanism of the effects of GSN on YTS cells, western blot was performed to detect the components of the PI3K/Akt pathway. As shown in Figure 3, AKT expression in three groups was not significantly different. Moreover, phosphorylation of Akt is characteristic of PI3K activation. The levels of PI3K and p-AKT of YTS-GSN cells were both significantly decreased, compared to YTS-Con cells. The results revealed that upregulation of GSN could inhibit the PI3K/Akt pathway.

**Figure 3.**
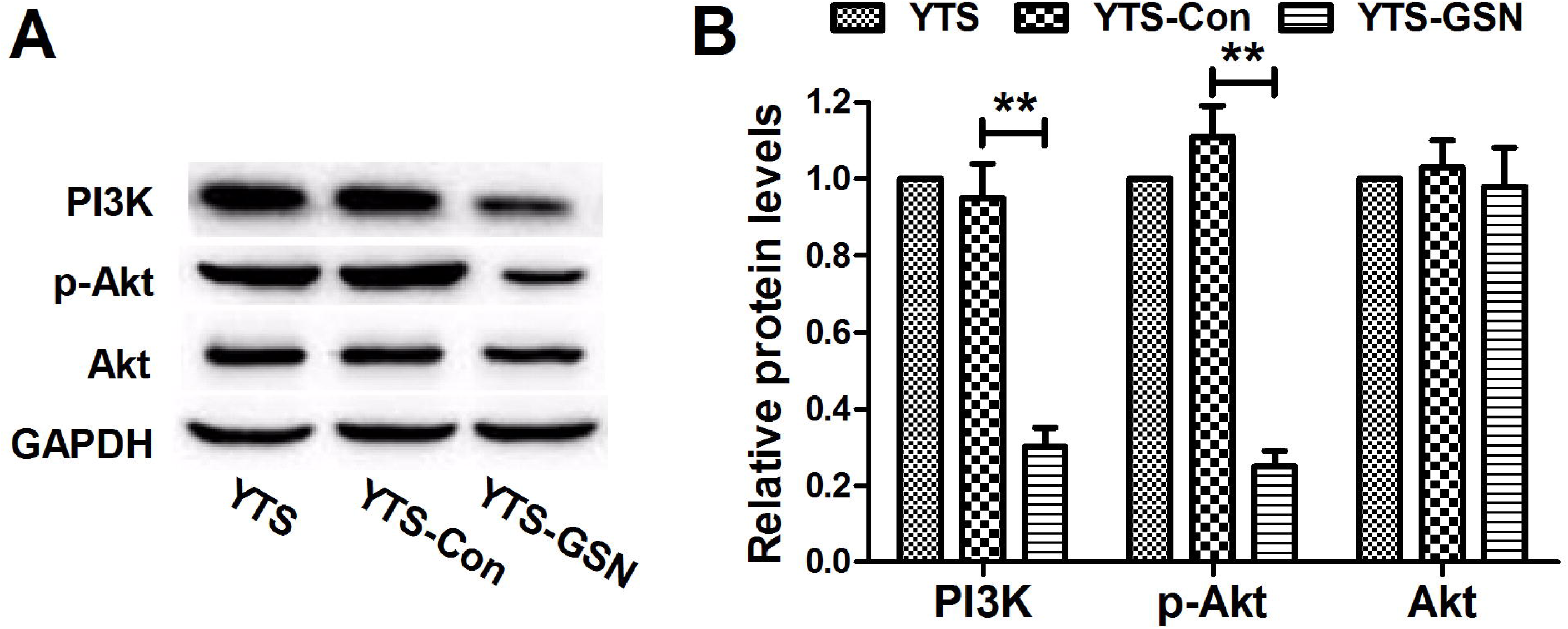
GSN overexpression inhibits the PI3K/AKT pathway in YTS cells. (A and B) Western blot analysis revealed that AKT expression in three groups was not significantly different. The protein levels of PI3K and p-AKT of YTS-GSN cells were significantly reduced at 48 h, compared to YTS-Con cells. \*\**P* < 0.01.

### Blocking the PI3k/AKT pathway inhibits the proliferation and invasion of YTS cells and promotes apoptosis

To confirm whether blocking the PI3K/Akt pathway inhibits cell proliferation and invasiveness and promotes apoptosis of YTS cells, LY-294002, a specific inhibitor of PI3k which can significantly inhibit the protein expression of p-AKT and PI3K but not AKT, was used to treat cells (Fig. 4A and 4B). As expected, CCK-8 assay and colony formation assay showed that blocking the PI3K-Akt pathway caused increased cell proliferation and colony formation (Fig. 4C and 4D). Flow cytometry analysis and transwell invasion assay exhibited that blocking the PI3K/Akt pathway resulted in an increased apoptosis and diminished cell invasion ability in YTS cells (Fig. 4E and 4F).

**Figure 4.**
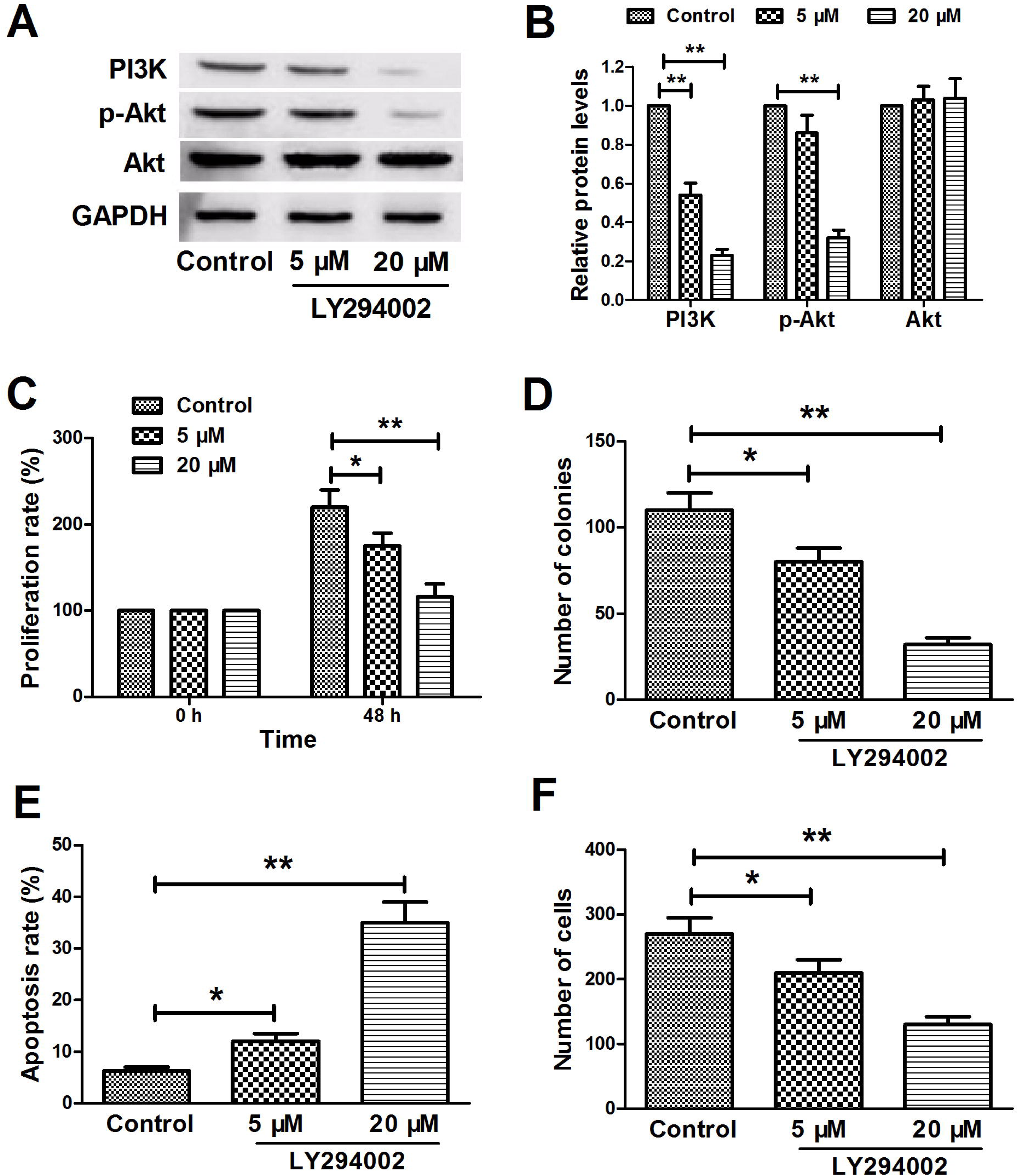
Blocking the PI3K/AKT pathway inhibits YTS cell proliferation and invasion and promotes apoptosis. (A and B) Western blot analysis revealed that LY294002 treatment (5 μM and 20 μM) significantly suppressed the p-AKT and PI3K in a dose-depended manner, while AKT expression in three groups was not significantly different. (C) CCK-8 assay showed that LY294002 treatment had an inhibitory effect on cell proliferation of YTS cells in a dose-depended manner at 48 h. (D) Colony formation assay showed that LY294002 treatment caused a decrease in the clonogenic survival of YTS cells in a dose-depended manner. (E) Flow cytometric analysis exhibited that LY294002 treatment had a promotive effect on cell proliferation of YTS cells in a dose-depended manner at 48 h. (F) Transwell invasion assay showed that LY294002 treatment had a inhibitory effect on cell invasion of YTS-GSN cells in a dose-depended manner at 48 h. \**P* < 0.05,\*\**P* < 0.01.

## Discussion

NKTCL is a common kind of malignant lymphoma, and usually develops in the nasal cavity but can also occur in extranasal sites, either as a primary extranasal or disseminated disease (Takeuchi et al., 2014). A recent study exhibits that the level of GSN is significantly decreased in serums of the advanced NKTCL patients. However, the potential effect of GSN on NK/T-cell lymphoma cells and molecular mechanism remain unclear.

GSN is a protein that is broadly expressed intracellularly, including in the cytoplasm and mitochondria and exists in both an intracellular and an extracellular form (Wen et al., 1996). Previous studies reveal that the expression level of GSN is decreased in many cancers, including NKTCL (Tanaka *et al*. 2006, Zhou *et al*. 2016). Deng *et al* show that the upregulation of GSN promotes cell growth and motility and speculate, which is involved in the progression of human oral cancers (Deng et al., 2015). Nevertheless, a study reveals that overexpression of GSN reduces the proliferation and invasion of colon carcinoma cells (Li et al., 2016). Our study revealed that GSN overexpression significantly suppressed cell proliferation and invasion in YTS cells. A previous study shows that GSN suppresses apoptosis by negatively regulating the expression of apoptosis-associated genes in hepatocarcinoma cells (Zhou *et al*. 2015). Our result showed that overexpression of GSN significantly increased apoptosis of YTS cells. Nevertheless, Abedini *et al* exhibit that GSN plays roles as both effecter and inhibitor of apoptosis, which underlines its association in a wide variety of cancer types (Abedini et al., 2014). According to the above results, GSN have different effects on cell proliferation, apoptosis and invasion in different cancers, which may be caused by GSN activating or inactivating different signaling pathways in varying cancers.

The PI3K/Akt pathway plays a vital role in cell survival by suppressing apoptosis and promoting cell proliferation (Vivanco and Sawyers 2002). Akt, an essential serine/threonine kinase is a crucial component of the PI3K signaling pathway whose activation has been involved in the genesis or progression of many human malignancies (Blume-Jensen and Hunter 2001, Vivanco and Sawyers 2002). Previous studies show that AKT1 and AKT2 that are the target genes of PI3K are overexpressed in breast, gastric and ovarian cancers (Staal 1987, Bellacosa et al., 1995). Many studies demonstrate that the constitutively active PI3K or Akt is oncogenic in cell systems and animal tumor models (Chang et al., 2003, Liu et al., 2015). Several studies have shown that Akt/PKB are involved in immune activation, cell proliferation, apoptosis and cell survival through activating the transcription of varieties of genes (Fowles et al., 2015, Warfel and Kraft 2015). Our study revealed that significantly upregulation of GSN inhibited the PI3K/AKT pathway in YTS cells. A previous study revealed that the cytoskeletal protein, GSN, was a vital determinant of cell invasion and scattering by inhibiting E-cadherin expression through the HGF-PI3K-Akt signaling pathway in gastric cancer (Huang et al., 2016). In addition, it is reported that constitutive PI3K/Akt activation promotes the progress of prostate cancer from organ-confined disease to highly invasive and even possibly metastatic disease. Due to its vital regulator of cell survival, Akt has considered as a crucial factor in tumorigenesis (Nowinski et al., 2015). Consistent with that, in our study, blocking the PI3K/Akt pathway inhibited cell proliferation and invasion of YTS cells, while promoted apoptosis.

## Conclusion

We speculated that GSN overexpression inhibited cell proliferation and invasion and promoted apoptosis of YTS cells, which was closely related to NKTCL and might have an anti-tumor effect. However, to our knowledge, relevant reports on the association between GSN and NKTCL are relatively few. Therefore, the specific pathogenesis requires further investigation.

## Materials and methods

### Cell lines and culture

The NK cell line YTS was purchased from ATCC (Manassas, VA) and maintained in RPMI 1640 medium supplemented with 10% FBS (Takara Biotechnology Co., Ltd., Dalian, China), 1% NEAA (Invitrogen, Carlsbad, CA, USA), 1% sodium pyruvate, 10 mM HEPES (PAA), 2 mM L-glutamine (Biochrom, Berlin, Germany), and 1% penicillin streptomycin (100 μg/ml; Invitrogen Life Technologies, Beijing, China) and 5% CO_2_ at 37°C.

The human embryonic kidney (HEK) 293T was purchased from the cell bank of the Chinese Academy of Sciences (Shanghai, China). The 293T cells were maintained in DMEM (Hyclone, Logan, UT) supplemented with 10% FBS, 10 mM HEPES, and 1% penicillin streptomycin and 5% CO_2_ at 37°C.

### Plasmids

The lentiviral vector we used was pCDH-CMV-MCS-EF1-copGFP (DCE; System Biosciences, Mountain View, CA) in this study. The packaging plasmids were pCMV-Δ8.2 and pCMV-VSV-G (System Biosciences). The GSN plasmid was purchased from Sino Biological Inc.

### Construction of the Lenti-GSN vector and lentivirus packaging

A specific primer was designed using Primer Premier 5.0 software (Shanghai Shenggong Biology Engineering Technology Service, Ltd., Shanghai, China) according to the nucleotide sequences of the human GSN gene, as reported in Genebank (http://www.ncbi.nlm.nih.gov/genbank/; reference sequence: NM_000177). The primer sequence for GSN was as follows: DCE-GSN-F: 5’-attctagagctagcgaattcATGGCTCCGCACCGCCCCG-3’; and DCE-GSN-R: 5’-ccttcgcggccgcggatccTCAGGCAGCCAGCTCAGCC-3’. The coding DNA sequence region of the GSN gene was amplified in the Perkin-Elmer (Gene Amp PCR system 2400) thermal cycler according to the manufacturer’s instructions. The target DNA gene fragment was subcloned into the DCE lentiviral vector to construct a GSN overexpression lentiviral vector (lenti GSN).

293T cells were cultured in 10 cm cell culture dishes (3 × 10^6^ cells/dish). The lentiviral vector packaging system made as the following: a solution of 500 μl was first prepared consisting of 12 μg of plasmid pCMV-Δ8.2, 10 μg of pCMV-VSV-G of, 22 μg of transfer expression plasmid lenti-GSN, and 125 μl of 2 mM CaCl_2_ in deionized distilled water. CaCl_2_/DNA was then added dropwise while vortexing to equal volume of 2 × HBS for a total of 1mL and was added to the cells at a density of 80%. The GFP expression was observed by fluorescent microscopy after 24h. The supernatant was harvested by centrifugation at 3000 rpm for 5 min at 4°C after 48h and the ratio of positive cells was measured by using FACSCalibur flow cytometer (Becton Dickinson, Franklin Lakes, NJ). The high-concentration lentiviral concentrate was used to infect the YTS cells.

### Lentiviral transfection of the YTS cells

YTS cells were seeded in 24 well plates (4 × 10^4^ cells/well). The viral supernatant with Lenti-GSN was added into the cells at a density of 70%-80%. After 72 h, the transfection ratio was determined under a fluorescence microscope and was measured by flow cytometry. The cells with a transfection ratio of >70% served as the target cells and were identified by qRT-PCR analysis.

### Real time quantitative polymerase chain reaction (RT-qPCR) analysis

Total RNA was isolated from cultured cells using TRIZOL reagent (Invitrogen, Carlsbad, CA). 2 μg total RNA was then reverse-transcribed using the Transcriptor First Strand cDNA synthesis Kit (Roche, Mannheim, Germany) with random hexamers. GSN mRNA was detected using Fast SYBR green PCR master mix (PE Applied Biosystems, Foster City, CA) according to the manufacturer’s protocol and the primer sequence for GSN and GAPDH was as follows: GSN-QF: 5’-GCT GAG GTT GCC GCT GGT G-3’, and GSN-QR: 5’-TGT GTT GGT TGC ATT TCC TTT TTG-3’; GAPDH-F: 5’-TGG TAT CGT GGA AGG ACT CAT GAC-3’, and GAPDH-R: 5’-ATG CCA GTG AGC TTC CCG TTC AGC-3’. Relative mRNA expression of GSN was calculated with the comparative threshold cycle (Ct) (2^−ΔΔCt^) method.

### Cell proliferation assay and soft agar colony formation assay

CCK-8 assay was performed to detect the growth of YTS, Lent-Con-transfected YTS, as well as Lent-GSN-transfected YTS cells. Cells were seed in 96-well plates at a density of 1 × 10^4^ cells/well and were incubated for 24, 48, 72 or 96 h in a humidified incubator. Subsequently, 10 μl CCK-8 solution (7Sea PharmTech, Shanghai, China) was added to the wells at indicated time. After incubation for 3-4 h, the absorbance was detected using the using a multilabel counter (Enspire Multimode Plate Reader, PerkinElmer, USA) at 450 nm.

For colony formation assay, all 6-well culture plates containing the bottom and soft layers were used. The cells were plated out in soft agarose as follows: cells were harvested from monolayer culture, washed and resuspended at 4 × 10^4^ cells/ ml in fully supplemented RPMI 1640 culture medium and molten 1.5% agarose (to a final concentration of 0.3%) on Day 0 A 0.5 ml of cellular suspension was applied to the base layer (1 × 10^4^ cells/well) and then allowed to set at 4°C for 6 min. Duplicate soft agarose cultures were established to assess colony formation. Cultures were placed in a incubator 37°C, 5% CO_2_ and 100% relative humidity for 12 days. The number of colonies containing ≥ 50 cells was counted using a light microscope.

### Cell apoptosis assay

Cell apoptosis was detected using the Annexin V-phycoerythrin (PE)/7-amino-actinomycin D (7-AAD) Apoptosis Detection Kit (Nanjing KeyGen Biotech Co., Ltd., Nanjing, China) according to the manufacturer’s instructions. Cells were seed in 6-well plates at 5 × 10^5^ per well. The cells were harvested and washed twice in PBS. 1 × 10^6^ cells were resuspended in 500 μl binding buffer. The suspension was stained with 1 μl Annexin V-PE in the dark for 10 min at room temperature. A 5 μl of 7-AAD was added to the suspension and maintained for 10 min at room temperature in the dark. Cell apoptosis was analyzed using the BD FACSAria II cell sorter (BD Biosciences, Franklin Lakes, NJ, USA).

### Transwell invasion assay

For invasion assays, transwell filters (Corning Incorporated, Corning, NY, USA) were coated with Matrigel (BD Biosciences) for 24 h. 2× 10^5^ cells were seed into the upper compartment of the chambers with 100 μl serum-free RPMI-1640 medium. The lower chamber of the transwell was filled with culture media containing 10% FBS as a chemo-attractant. After 48 h incubation, non-invaded cells on the top of the transwell were scraped off with a cotton swab. Cells successfully translocated were fixed with 10% formalin and counted under a light microscope.

### Western blot analysis

Total protein was extracted from cells using lysis buffer (Roche Diagnostics, Basel, Switzerland). Protein samples (30 μg) were separated by 10% SDS-PAGE and transferred onto polyvinylidene fluoride (PVDF) ultrafiltration membrane (Sigma-Aldrich, Corp., Cambridge, UK) for 2 h at 4°C. The membranes were blocked with 5% non-fat milk for 1 h at room temperature. The membranes were washed three times for 5 min each with 15 ml TBS Tween 20 (TBST; Cell Signaling Technology, Inc.). The membranes were incubated with primary antibodies overnight at 4°C. The membranes were then incubated with horseradish peroxidase-conjugated antibody for 2 h at 37°C. Antigen-antibody complexes were visualized by enhanced chemiluminescence (ECL) blotting analysis system (Amersham Pharmacia Biotech, Buckinghamshire, UK) and GAPDH served as the internal reference. The primary antibodies used in this study are as follows: GSN, PI3K, AKT, p-AKT, and GAPDH antibody (1:1,000; Cell Signaling Technology, Danvers, USA). An HRP-conjugated anti-rabbit IgG antibody was used as the secondary antibody (Santa Cruz, Santa Cruz Biotechnology, USA).

### Statistical analysis

The data are presented as the mean ± SD and analyzed using *t* test or ANOVA by SPSS Software version 19 (SPSS, Inc., Chicago, IL, USA). *P* < 0.05 was considered to be statistically significant difference.

### Author contributions

Yanwei Guo, Hongqiao Zhang and Mingzhi Zhang conceived and designed the experiments; Xin Xing, Lijuan Wang, Jian Zhang Lin Yan and Xiaoke Zheng performed the experiments and participated in data analyses; Yanwei Guo and Xin Xing wrote the manuscript.

